# Inferring lifestyle for Aves and Theropoda: a model based on curvatures of extant avian ungual bones

**DOI:** 10.1101/517375

**Authors:** Savannah E. Cobb, William I. Sellers

## Abstract

Claws are involved in a number of behaviours including locomotion and prey capture, and as a result animals evolve claw morphologies that enable these functions. Past authors have found geometry of the keratinous sheath of the claw to correlate with mode of life for extant birds and squamates; this relationship has frequently been cited to infer lifestyles for Mesozoic theropods including *Archaeopteryx.* However, claw sheaths rarely fossilise and are prone to deformation; past inferences are thus compromised. As the ungual phalanx within the claw is relatively resistant to deformation and more commonly preserved in the fossil record, geometry of this bone would provide a more useful metric for paleontological analysis. In this study, ungual bones of 108 birds and 5 squamates were imaged using X-ray techniques and a relationship was found between curvatures of the ungual bone within the claw of pedal digit III and four modes of life; ground-dwelling, perching, predatory, and scansorial; using linear discriminant analysis with Kappa equal to 0.69. Our model predicts arboreal lifestyles for certain key taxa *Archaeopteryx* and *Microraptor* and a predatory ecology for *Confuciusornis.* These findings demonstrate the utility of our model in answering questions of palaeoecology, the theropod-bird transition, and the evolution of avian flight.

## Introduction

The amniote claw is utilised in multiple functions related to ecology and lifestyle. Claws bear an animal’s weight during locomotion, are utilised in prey capture, and more. These tasks exert selective pressure on the claw and so claws are expected to evolve morphologies that enable performance of essential functions whilst minimising stress/strain during locomotion^1,2^. The evolutionary reduction of bending stresses during terrestrial locomotion is the proposed cause of the relatively flat claws observed for ground-dwelling taxa compared to pedal claws belonging to arboreal and/or predatory taxa, which tend to possess more curved claws for enabling grip^1-10^. The relationship between claw morphology and lifestyle has frequently been utilised to infer lifestyle for fossil taxa.

Curvature of extant claws as quantified using “claw angle”, or the angle found for the arc of a circle approximated using points on the claw^11^, has been shown to correlate with lifestyle for a diverse group including avians, squamates, and mammals^3-8^. Though curvature and other aspects of external claw geometry (i.e. measures taken on sheaths, toe pads, and skin) are known to correlate with lifestyle and locomotor ability^4,6,7,8,12,13^, internal structure of the claw has not been sufficiently studied to this purpose. Amniote claws are comprised of a terminal phalanx encompassed by an external keratinous sheath^12,14-19^.

For avian and reptilian claws, these terminal phalanges are known as ungual bones, and the external sheaths are comprised of rigid, insoluble β-keratins and ‘soft’ α-keratins. The filamentous structure of keratins causes viscoelastic behaviour when hydrated^20^, and so claw sheaths are likely more prone to taphonomic distortion than are internal ungual bones. Even well-preserved claw sheaths may have variable claw angles due to wear during life, whereas ungual bones experience no abrasive contact with the substrate and are thus unaffected by wear. Using Extant Phylogenetic Bracketing (EPB)^21^, we infer non-avian theropod dinosaurs and other fossil taxa within Theropoda and Aves would have possessed claws of similar composition with comparable material properties^20,22,23^. If so, then it seems likely fossil taxa on the avian lineage evolved similar claw morphologies to extant birds in response to similar ecological and functional pressures.

Theropod claw sheaths are rare but not unknown from the fossil record^24-33^. However, for most fossil specimens, the claw sheath is either broken or entirely absent leaving only the ungual bone, and fossilised toe pads or skin are even rarer^34-43^. Measurements of fossil claw angle tend to be either based on reconstructions^4^ or are taken directly on ungual bones^5^ in past analyses. Past attempts to find a relationship between curvature of the keratinous sheath and the ungual bone for reconstruction purposes have yielded variable results^4,8,14,44,45^. Reconstructed claw angles are thus based on either supposition or on a relationship for which no consensus has been reached.

For these reasons, we believe predictive models based solely on extant sheaths are inherently flawed. Previous mode of life predictions for fossil taxa are thus compromised. One way to overcome the limitations of past studies is to study the ungual bone so that fossil claws not preserving soft tissues can be directly compared to extant claws. However, very little is known about avian or squamate ungual bone morphology.

As the ungual bone fits within the sheath, we expect a similar relationship exists between ungual bone morphology and mode of life as has been found for claw sheaths. This study investigates the relationship between dorsal and ventral curvatures of ungual bones and behavioural categories terrestrial, perching, predatory, and scansorial for a diverse group of extant avians and squamates. The results are then utilised to infer lifestyle for a sample of fossil paravians and avians.

## METHODS

We examined curvatures for ungual bones and sheaths belonging to 95 species of bird representing 25 orders and 5 species of squamate. As this study seeks to infer modes of life for fossil taxa on the avian lineage, the final predictive model is based on bird claws. A small group of behaviourally diverse squamates have also been tested to investigate if determined trends are universal or constrained by clade. Crocodilians have been excluded because modern taxa lack the behavioural diversity this study seeks to analyse.

Each specimen was placed into one of four behavioural categories; terrestrial, perching, predatory, or scansorial; based on the literature^46-58^. Claws from predatory and perching taxa possess similar ranges of curvature, and so predatory taxa were included to account for a potential confounding factor. Specimens have been selected such that each behavioural category includes three or more orders and some orders are represented within multiple behavioural categories. All extant specimens measured are adults to constrain potential influences of ontogenetic and/or behavioural changes during life known for some taxa^59^.

Claws were radiographed in lateral view using the Nomad Pro Radiography Unit (https://www.dentsplysirona.com/en) and processed in the SIDEXIS software (https://www.dentsplysirona.com/en). For small claws less than roughly 10 mm long, sheath data were supplemented with photographs and multiple images at various exposures were layered to create a weighted average in Helicon Focus 7 to improve resolution.

The Nomad Pro Radiography Unit has a lithium polymer battery that charges at 110/220 V and operates at 22.2 V and 1.25 Ahr. The anode voltage is 60 kV true DC and the anode current is 2.5 mA. We utilised a size 2 digital sensor with an active sensor area of 25.6 × 36 mm and external dimensions of 31.2 × 43.9 × 6.3 mm. The system acquires images with a measured resolution of 16 Lp/mm, and maximum dose of radiation to the user registered between 0.0117 mGy and 0.0310 mGy at the palm depending on how the device is positioned^61^. All measured dosages are well below permissible limits according to the Ionising Radiation Regulations 2017^61^.

This device enabled the rapid, inexpensive, on-site acquisition of a large data set and is a practical alternative to CT scanning. However, more refined techniques such as geometric morphometric (GM) analysis may not be feasible based on pixel count and so linear measurements have been taken and analysed.

Photographs of fossil specimens were acquired from published sources (details in Table S4). Fossil specimens include paravians, avialans, and avians and were selected based on condition, image resolution, and phylogenetic closeness to extant birds. The selected fossil ungual bones show no obvious signs of breakage or distortion, and so we assume values measured for fossil ungual bones represent true claw angles of the animal during life. For fossil claws in possession of keratinous sheaths, some level of degradation is apparent for all specimens and slight reconstruction was necessary.

Claw length was limited to a maximum of 44 mm to fit on the active sensor area and a minimum of 7 mm because fine details could not be resolved for very small claws. As body mass and claw radius correlate^8^, body masses for the sample taxa are limited from 36g to 1930g to constrain claw size.

Body masses were determined from the CRC Handbook of Avian Body Masses^62^ and may differ slightly from true body masses of individuals as this information was often unavailable for the sample specimens. When it was not possible to sex the specimens, body mass was calculated as the average of male and female body mass. We regressed body mass against claw angle for each behavioural category to determine effects of scaling.

Most sampled claws are museum specimens that were imaged on-site. The dataset was also supplemented with specimens were acquired from independent sources. Full specifications of the specimens are listed in Table S3.

Claw curvature between digits can vary significantly^14^ and so we have constrained digit of study to pedal digit III after the fashion of past studies. Digit III is the longest digit for many birds, squamates, and non-avialan Mesozoic theropods and represents the first and last point of contact with the substrate during terrestrial locomotion. As it is functionally significant, claw morphology of digit III is expected to be well-adapted for locomotion and other functions^6,63^.

Due to phylogenetic conflict within Aves and the difficulties in placing fossil taxa phylogenetic corrective methods were not used. Two recent studies resolved conflicting topologies for Aves^64,65^, and phylogenies for fossil taxa are less stable.

## Geometric measurements

Angles of curvature for dorsal or ventral surfaces of each claw were calculated using three landmark points on the claw (Figure 1). Ventral and dorsal curvatures of the ungual bones (IU, OU) and claw sheaths (IS, OS) were measured according to the methods outlined in Feduccia (1993) and Pike and Maitland (2004). Three points A, B, and X are placed on each claw for calculating angle of curvature (see Figures 1 and 2). Custom software - DinoLino.exe - was created in Microsoft Visual Studio using C# for taking these measurements with improved speed and precision as compared to other available programs (https://github.com/johnwelter/Dino-Lino).

**Figure 1.**
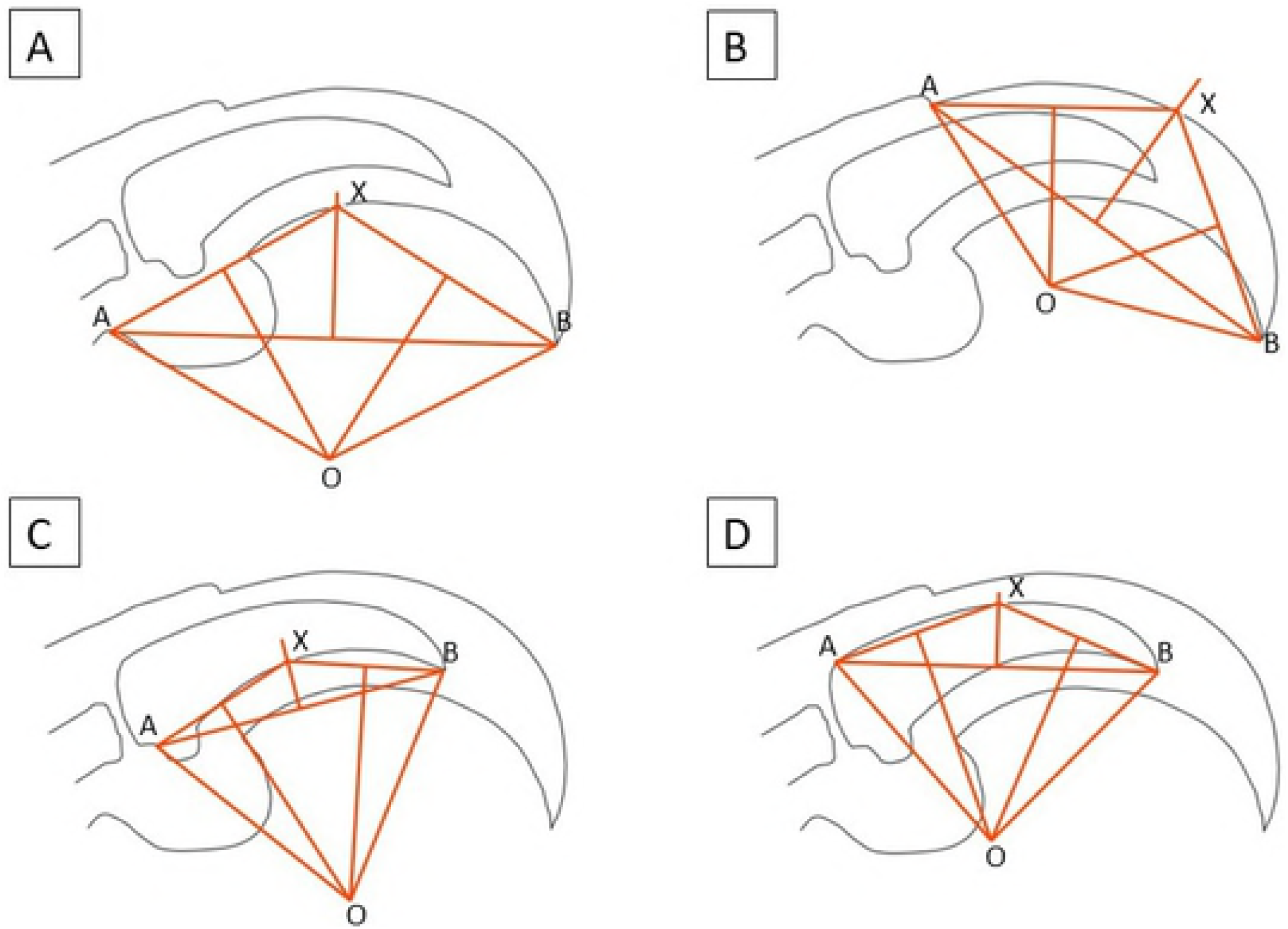
Methods of determining claw angle. Claw drawn in Inkscape; geometric measurements performed in DinoLino.exe. A. Feduccia’s method (1993) of quantifying claw angle for inner curvature of the claw sheath, here denoted IS. B. Pike and Maitland’s method (2004) of quantifying claw angle for outer curvature of the claw sheath, here denoted OS. C. A modification of Feduccia’s method to measure inner curvature of the ungual bone, here denoted IU. Landmarks are placed at the base of the flexor tubercle (A), the tip of the ungual phalanx (B), and the intersection of 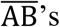 bisector with the underside of the ungual phalanx (X). The centre of a circle (O) is drawn at the intersection of perpendicular bisectors to 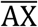 and 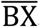, and claw angle is taken as ∠AOB^67^. D. A modification of Pike and Maitland’s method to measure outer curvature of the ungual bone, here denoted OU. Landmarks are placed at the proximal dorsal end of the ungual phalanx (A), the tip of the ungual phalanx (B), and the intersection of 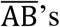 bisector with the underside of the ungual phalanx (X). Claw angle then follows the approach described in C.

**Figure 2.**
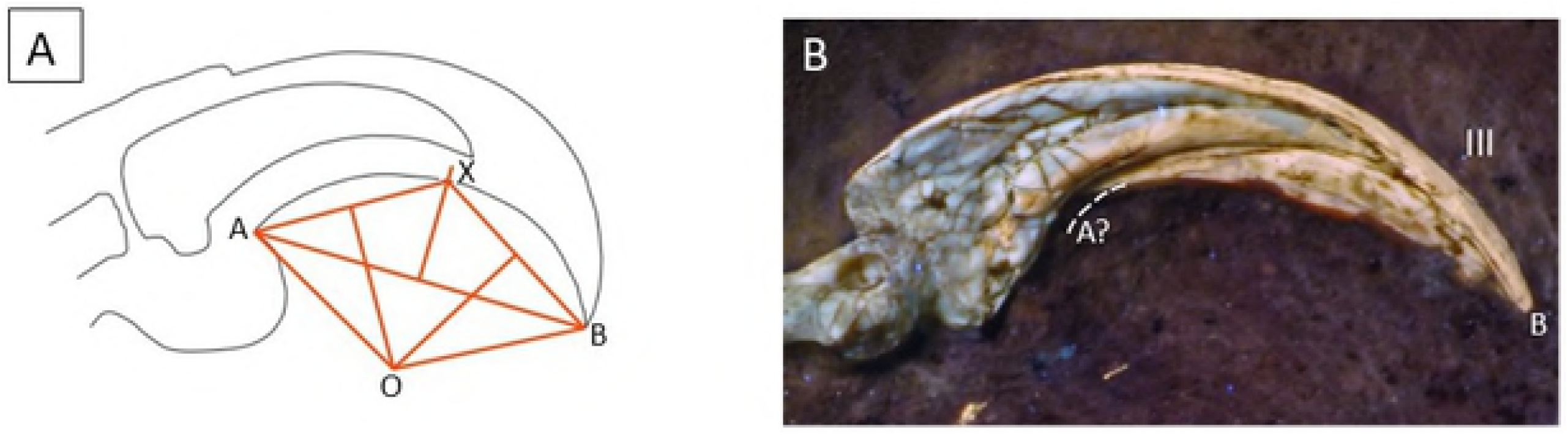
Alternative measure utilised for fossil claws and example *Archaeopteryx* claw. Claw drawn in Inkscape; measurements performed in DinoLino.exe. A. method of measuring claw angle^11^, here denoted IS2. Landmark A is located at the interface where ventral claw sheath meets toe pad. Landmark B is located at the sheath tip. B. Right pedal claw from the 12^th^ Archaeopteryx specimen^28^. Landmarks have been approximated using the method shown in A. Slight reconstruction was necessary to account for decay at the base of the sheath.

Two measures (IU, OU) were taken on fossilised unguals. For fossil claws in possession of keratinous sheaths, ventral curvature of the sheath was measured using a method excluding the toe pad that has also been shown to correlate with modes of life^3^ (Figure 2). This measurement shifts landmark A distally to the start of the dorsal sheath but retains landmark B at the tip of the claw sheath as this can often be directly measured on the fossil claws (Figure 2). Toe pads are not known for any of the measured fossil claws, and so using this measurement as opposed to IS minimises the amount of reconstruction necessary (Figure 2).

## Statistical analysis

Statistical analyses and graphical summaries were performed in R^67^. Body mass has previously been shown to have a complex relationship with claw morphology^8^, and so we tested for a linear relationship between body mass and claw angle by group for each measure taken on avian ungual bones and claw sheaths. Simple linear regression was performed by group on body mass and claw angles for the dataset of extant avians using the smatr package^68^. To normalise the data, body mass was log-transformed and claw angle was transformed by cosine. As variances differ among behavioural groups (heteroscedasticity), pairwise and non-parametric median and permutational tests were utilised to determine if median claw angles and/or centroids of combined measures of claw angle differ between groups. The statistically significant p-value was defined as the standard 0.05.

Measures were submitted to linear discriminant analyses using the caret package in R^69^. Predictive models were created using four subsets of the avian data: ungual bone measurements (IU, OU), claw sheath measurements (IS, OS), sheath and bone measurements (IU, OU, IS, OS), and sheath and bone measurements with the modified measure IS2. Predictive success of the models was tested for extant birds using bootstrap resampling with 2000 iterations using the train function of the caret package^69^. Predictions were generated for fossil taxa using the determined models.

## RESULTS

### Relationship to body mass

Figure 3 clearly shows that very little of the variation in claw angle can be explained by body mass although there is a weak association with outer curvature of the keratinous sheath. Ungual bones do not show a statistically significant relationship with body mass for the sample taxa. There is a negative relationship between dorsal claw angle of the sheath (OS) and body mass for terrestrial taxa (p=0.04641). However, the amount of data explained by the linear best fit line between OS angles and body masses of terrestrial taxa was low (R^2^=0.1219), suggesting other factors are influencing this relationship.

**Figure 3.**
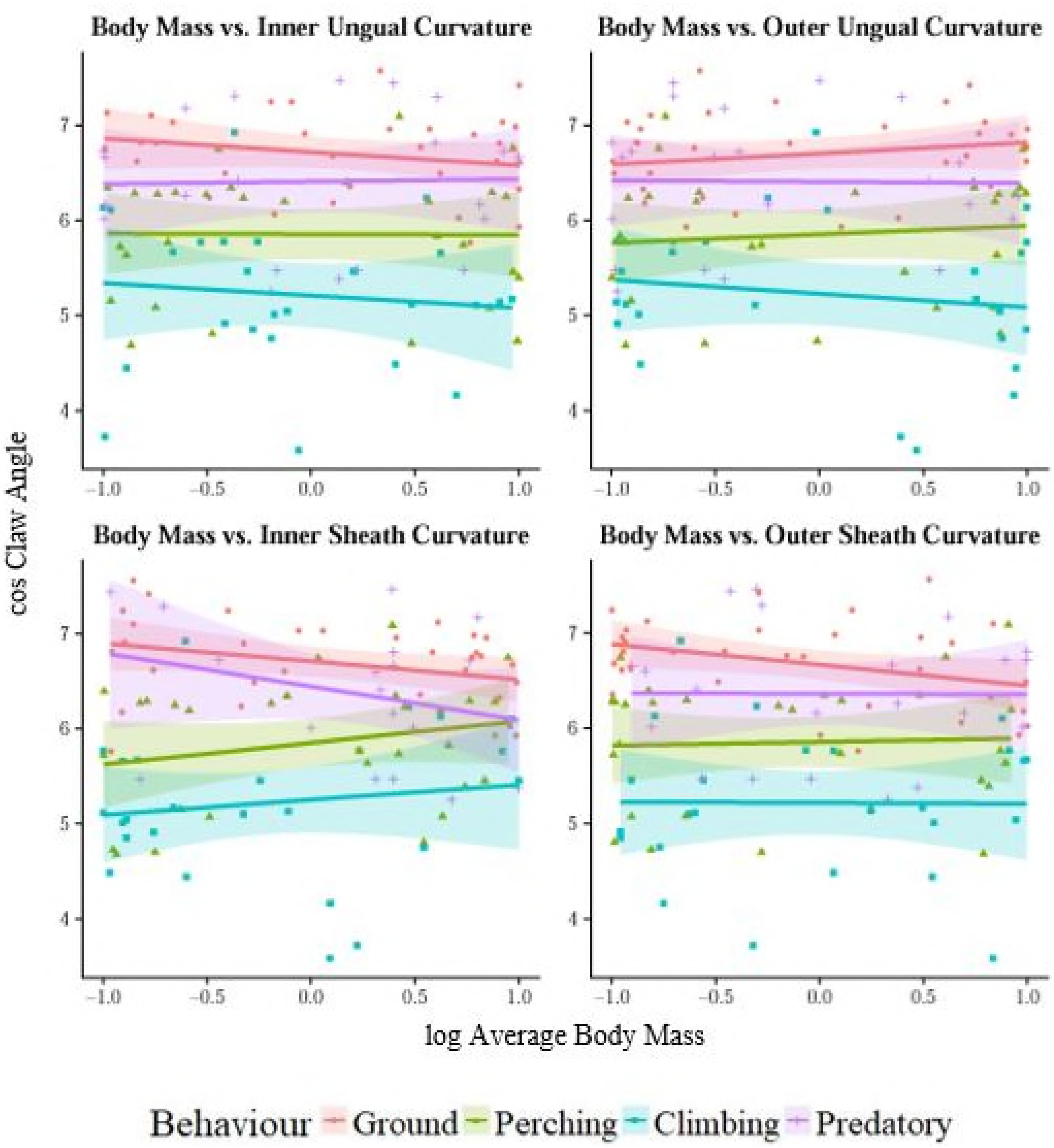
Regression plots for claw angle against body mass grouped by behavioural category. The shaded area represents 95% confidence limits of the scatter.

### Claw geometry and behavioural category

The relationship found for ungual bones is similar to that found for claw sheaths: lower angles of curvature correlate with terrestrial lifestyles, intermediate claw angles correlate with perching and predatory lifestyles, and higher claw angles correlate with scansorial lifestyles. Compared to claw sheaths, ungual bone curvatures had relatively constrained ranges and possessed lower values of claw angle. Perching, ground-dwelling, and climbing taxa had roughly equivalently sized ranges of ungual bone curvatures. Predatory taxa had the smallest ranges of claw angle but, notably, were represented by the fewest sample specimens. Median values of inner curvature were significantly lower than those of outer curvature in all but the ‘predatory’ group.

Boxplots for inner (A) and outer (B) claw curvatures distinguished by behavioural category for all extant taxa. Shaded boxes depict interquartile range (IQR) and whiskers depict distance between IQR and points up to 1.5 distances from the IQR. Outliers are represented with circles or stars dependent on taxonomic group and are greater than 1.5 distances from the IQR. Morphospaces based on LD1 and LD2 generated by an LDA of combined ungual bone measures (C) and combined sheath and bone measures (D) for extant birds, overlain with data for fossil claws. Ellipses were drawn with 95% confidence from the centroid for each group.

All squamate claws plotted as outliers and have been excluded from further analyses for their likely incomparability with extant avians and fossil theropods. Ranges for measures taken on claw sheaths, particularly inner sheath curvatures, showed more outliers than those measured for ungual bones, suggesting high variability of ungual soft tissues relative to bone. Of the sampled avian ungual bones, extreme values are apparent in the ‘terrestrial’ category showing inner curvatures measured at zero: *Numenius arquata* (Eurasian curlew), *Larus canus* (common gull), and *Lagopus lagopus* (willow ptarmigan). Gulls and curlews have aquatic habits and webbed feet, which may have influenced the evolution of very flat ungual bones, and the willow ptarmigan has a visibly unusual claw morphology (Figure 5). Avian claws display a wide range of morphologies and so we believe these nearly flat ungual bones, though plotted here as extreme values, represent normal diversity in the population. The above-mentioned avian taxa have thus been retained in the analysis.

**Figure 4.**
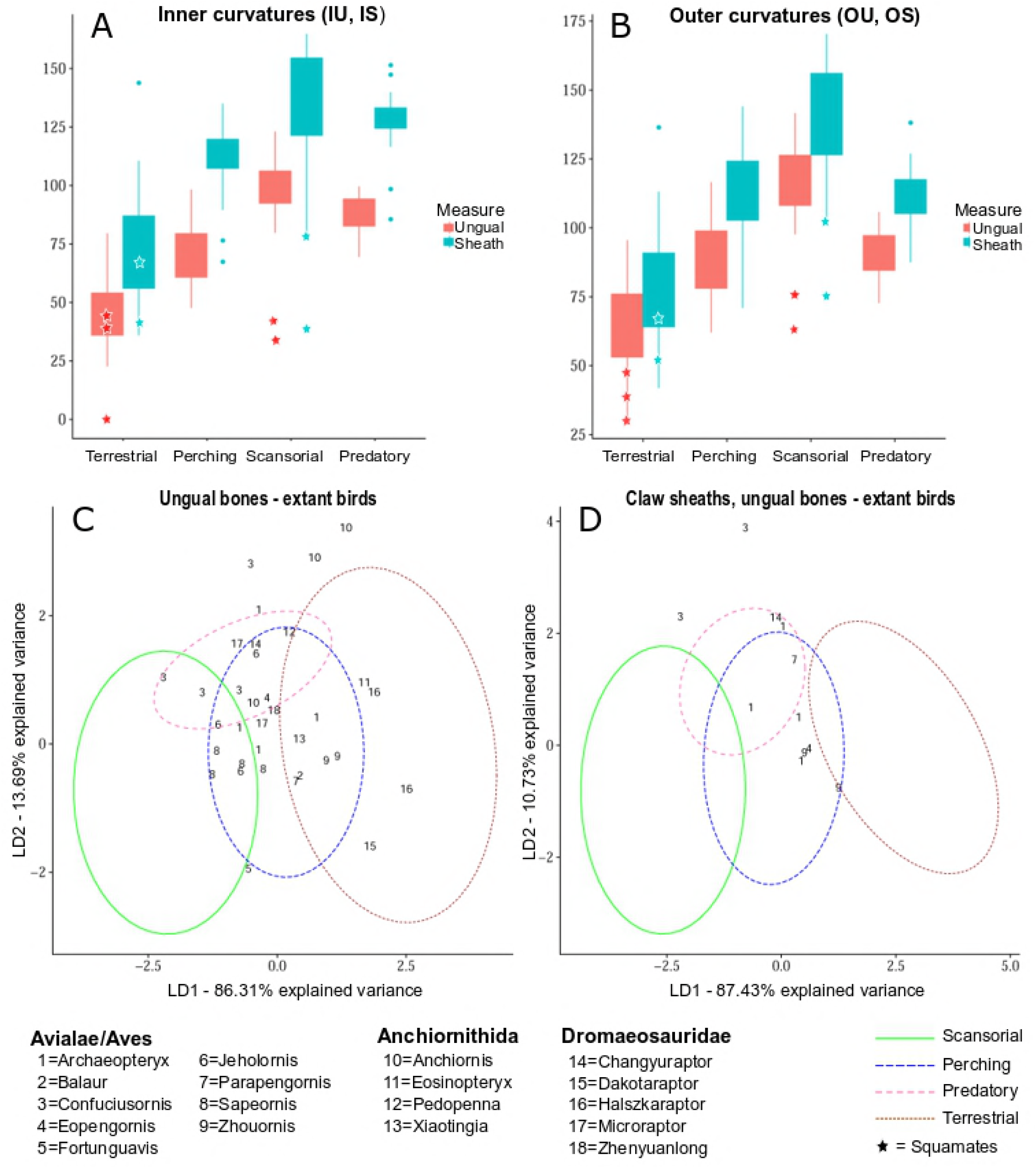
Curvatures of ungual bones and claw sheaths, digit III, for all extant and fossil taxa.

**Figure 5.**
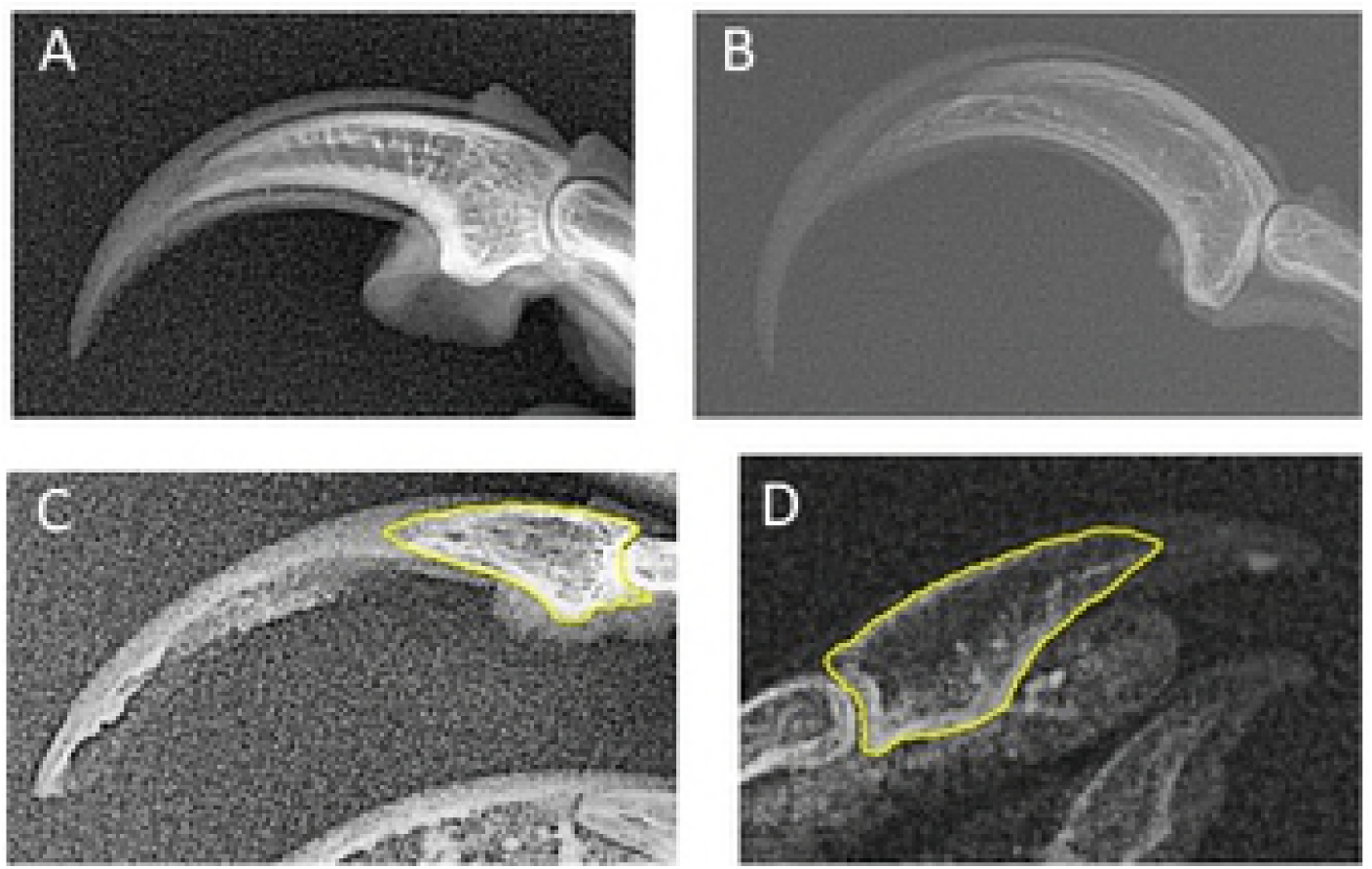
Radiographs of pedal digit III claws exhibiting significant morphological disparity. Ungual bones outlined in yellow have inner curvatures measured at zero. A. Left claw of kakapo (*Strigops habroptila*). Unregistered skin, National Museum of Scotland B. Left claw of ivory-billed woodpecker (*Campephilus principalis*). Unregistered skin, National Museum of Scotland. C. Right claw of willow ptarmigan (*Lagopus lagopus*). Skin specimen 1984.2.197, Liverpool World Museum. D. Left claw of gull (*Larus canus*). Skin specimen 1384, National Museum of Scotland.

For the model based solely on two ungual bone measurements (Figure 4, C), two axes LD1 and LD2 are generated that account for all variation. For the model utilising four measurements (Figure 4, D), LD1 and LD2 account for 97% of the variation, and so LD3 is not likely to significantly impact separation. LD1 represents greater or lesser values of claw angle for any given metric, accounts for greater than 85% of variation, and separates terrestrial taxa from predatory/perching taxa from scansorial taxa. LD2 represents the relationship between inner and outer curvatures of the claw, with higher LD2 values signifying a relatively high ratio of inner to outer curvature, and vice versa, and separates the predatory morphospace from the perching morphospace. Claws of perching birds tended to have lower LD2 values compared to claws of predatory birds, which have, on average, IU:OU roughly equal to one resulting in a slender claw with little tapering.

Despite differing medians (Table 2), there is overlap between ranges for each group. Morphospaces created based on measures of avian ungual bones overlap between all categories except ‘terrestrial’ and ‘scansorial’. Those based on all claw measures have similar degrees of overlap but manage to also separate ‘terrestrial’ and ‘predatory’ groups.

**Table 1.** Summary from a simple linear regression on cos(claw angle) against log(average body mass).

| Curvature | Behaviour | Slope | Intercept | p-value | R<sup>2</sup> |
| --- | --- | --- | --- | --- | --- |
| Inner Ungual | Terrestrial | -0.3324 | 2.4224 | 0.2277 | 0.04658 |
|  | Perching | -0.009344 | 0.021829 | 0.9658 | 6.94E-05 |
|  | Predatory | 0.02547 | -0.11135 | 0.9117 | 0.00063 |
|  | Scansorial | -0.0817 | 0.3474 | 0.6211 | 0.0108 |
| Outer Ungual | Terrestrial | 0.3062 | -2.153 | 0.2983 | 0.03484 |
|  | Perching | 0.1257 | -0.7192 | 0.58 | 0.01149 |
|  | Predatory | -0.01407 | -0.13774 | 0.95 | 0.000202 |
|  | Scansorial | -0.1542 | 0.8701 | 0.4757 | 0.02235 |
| Inner Sheath | Terrestrial | -0.5287 | 3.6631 | 0.07161 | 0.1009 |
|  | Perching | 0.2348 | -1.3762 | 0.2271 | 0.05354 |
|  | Predatory | -0.2106 | 1.5779 | 0.2314 | 0.07446 |
|  | Scansorial | 0.1219 | -0.8576 | 0.5081 | 0.01927 |
| Outer Sheath | Terrestrial | -0.5597 | 3.6567 | 0.04641 | 0.1219 |
|  | Perching | 0.04641 | -0.4269 | 0.8295 | 0.001748 |
|  | Predatory | -0.005653 | 0.112745 | 0.9793 | 0.00069 |
|  | Scansorial | -0.007677 | 0.009928 | 0.9681 | 7.11E-05 |

**Table 2.** Results of non-parametric tests performed on subsets of the avian data. P-values represent Bonferroni-corrected values.

|  | Wilcoxon Rank Sum |  |  | PERMANOVA |  |  |
| --- | --- | --- | --- | --- | --- | --- |
|  | Inner ungual curvature (IU) |  |  | Ungual curvatures (IU, OU) |  |  |
|  | Climbing | Ground | Perching | Climbing | Ground | Perching |
| Ground | 1.07E-09 | - | - | 0.0006 | - | - |
| Perching | 1.30E-10 | 4.01E-06 | - | 0.0006 | 0.0006 | - |
| Predatory | 7.13E-03 | 1.25E-07 | 4.29E-4 | 0.0006 | 0.0006 | 0.0036 |
|  | Outer ungual curvature (OU) |  |  | All measures (IU, OU, IS2, OS) |  |  |
| Ground | 1.07E-09 | - | - | 0.0006 | - | - |
| Perching | 5.60E-08 | 2.84E-06 | - | 0.0006 | 0.0006 | - |
| Predatory | 1.36E-06 | 1.48E-05 | 1 | 0.0006 | 0.0006 | 0.0736 |

Medians of inner and outer curvatures for ungual bones differed between all behavioural categories except the ‘perching’ and ‘predatory’ categories (Table 2). This is not unexpected as box plots shows inner and outer curvature values of the ‘predatory’ group completely overlap with ranges found for the ‘perching’ group (Figure 4 A, B). Pairing inner and outer curvatures of the ungual bone makes it possible to distinguish between perching and predatory claws; centroids based on these values differed significantly between all groups. Interestingly, including soft tissue measures in the multivariate analysis worsened separation between groups. Centroids based on four measures of sheath and ungual curvatures differed significantly between all groups except for perching and predatory taxa, the comparison of which yielded a p-value equal to 0.0736 after Bonferroni correction.

Kappa and average success are both reported; however, kappa is recommended when evaluating results because this metric accounts for unequal sample size by category. Accuracy by category varies, and total accuracy is influenced by unequal sample size of each behavioural group.

Predictive models based on ungual bone curvatures had greater success at classing extant bird claws (total accuracy = 0.7865, kappa = 0.7141) relative to models based on similar measures taken on keratinous sheaths (total accuracy = 0.7273, kappa = 0.6324). Both predictive models based on bone and sheath measurements had relatively greater success than models based on only two measures (Table 3). Utilising the modified sheath measure IS2 as opposed to IS slightly reduced predictive success (total accuracy = 0.8103, kappa = 0.7457) but did not greatly alter results relative to the model utilising IU, OU, IS, and OS (total accuracy = 0.8190, kappa = 0.7566).

**Table 3.** Predictive success of the models based on extant bird claws. Accuracy for each behavioural category, total accuracy, and kappa are listed.

| Included Measures | Accuracy: Terrestrial | Accuracy: Perching | Accuracy: Scansorial | Accuracy: Predatory | Accuracy: Total | Kappa |
| --- | --- | --- | --- | --- | --- | --- |
| Unguals (IU, OU) | 0.8182 | 0.658 | 0.813 | 0.8906 | 0.7865 | 0.7141 |
| Unguals, sheaths (IU, OU, IS, OS) | 0.8078 | 0.7948 | 0.8276 | 0.8594 | 0.819 | 0.7566 |
| Unguals, sheaths (IU, OU, IS2, OS) | 0.8279 | 0.7138 | 0.8347 | 0.8854 | 0.8103 | 0.7457 |
| Sheaths (IS, OS) | 0.7915 | 0.6506 | 0.6667 | 0.8083 | 0.7273 | 0.6324 |

**Table 4.**
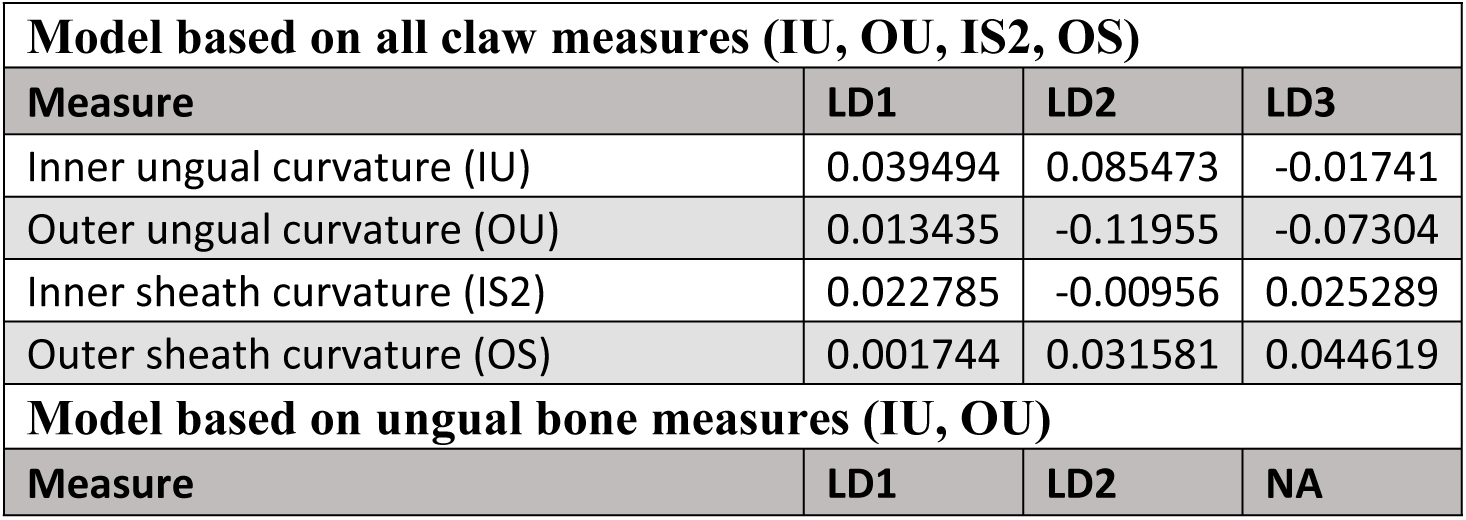
Variance-covariance loadings for each variable included in the analysis A. Variance matrix based on LDA of all measures B. Variance matrix based on LDA of ungual bones

| Model based on all claw measures (IU, OU, IS2, OS) |  |  |  |
| --- | --- | --- | --- |
| Measure | LD1 | LD2 | LD3 |
| Inner ungual curvature (IU) | 0.039494 | 0.085473 | -0.01741 |
| Outer ungual curvature (OU) | 0.013435 | -0.11955 | -0.07304 |
| Inner sheath curvature (IS2) | 0.022785 | -0.00956 | 0.025289 |
| Outer sheath curvature (OS) | 0.001744 | 0.031581 | 0.044619 |
| Model based on ungual bone measures (IU, OU) |  |  |  |
| Measure | LD1 | LD2 | NA |
| Inner ungual curvature (IU) | 0.041889 | 0.082856 | NA |
| Outer ungual curvature (OU) | 0.033374 | -0.09416 | NA |

Perching birds are consistently classed with the lowest accuracies ranging from 0.6506 to 0.7948 predictive success, and predatory taxa are classed with the highest accuracies ranging from 0.8083 to 0.8906 predictive success dependent on the dataset. Accuracies when classing terrestrial and scansorial taxa exceed 80% and were of similar values for the models based on all subsets except that based on sheaths, for which accuracy when classing terrestrial taxa (0.7915) was significantly greater than accuracy when classing scansorial taxa (0.6667).

### Comparison with fossil taxa

For both morphospace plots based either on ungual bone data or all claw measures, the majority of fossil taxa plot in the predatory/perching morphospaces. Dromaeosaurid and anchiornithid ungual bones plot in predatory, perching, and terrestrial morphospaces, and avialan ungual bones plot in predatory, perching, and scansorial morphospaces. For both morphospace plots based either on ungual bone data or all claw measures, the majority of fossil claws plot within the overlapping regions between 95% confidence ellipses, but some taxa plot in regions distinct to a particular morphospace. These include fossil dromaeosaurids *Dakotaraptor* and *Halszkaraptor* and anchiornithid *Eosinopteryx*, which all plot as terrestrial, and dromaeosaurid *Zhenyuanlong*, which plots as perching. No fossil claw plots as distinctly scansorial though some avialan claws including *Sapeornis, Confuciusornis*, and *Fortunguavis* plot in the shared spaces between predatory-scansorial and perching-scansorial taxa.

In the morphospace plot based on ungual bone data, fossil claws representing two *Anchiornis* and one *Confuciusornis* specimen represent outliers. One *Archaeopteryx* claw lies just outside the ‘predatory’ morphospace, but when measures of the sheath are included in the morphospace analysis, this data point shifts such that it is encompassed by the ‘predatory’ morphospace. In the morphospace plot based on all claw data, two *Confuciusornis* claws including the outlying specimen from Figure 4, C plot as outliers. It is unknown whether the outlying *Anchiornis* claws from Figure 4, C would be consistent outliers in both graphs as sheath data were unavailable for the measured specimens.

Claws belonging to *Archaeopteryx, Confuciusornis*, and *Microraptor* had a wide spread. *Archaeopteryx* specimens plotted within all morphospaces, and, interestingly, right and left digit III claws of the same specimen plotted quite far apart from each other. This was also the case for *Microraptor*, for which two claws belonging to a single specimen plotted in different morphospaces (predatory and perching) and received different classifications accordingly (Table S1). Though all were classed as predatory (Tables S1, S2), the four *Confuciusornis* claws were recovered in the overlaps between predatory, perching, and scansorial morphospaces and, in some instances, plotted outside of extant morphospaces altogether.

## DISCUSSION

### Utility of the model

Many previous studies have attempted to predict fossil lifestyles based on claw geometry, often utilising trends found for extant claw sheaths to classify fossil ungual bones. This study found a similar relationship between lifestyle and curvature for ungual bones as has been found for claw sheaths, but ungual bones possess relatively lower curvatures compared to sheaths (Figure 4 A, B) and thus cannot be directly compared to extant sheaths without risking misclassification. By determining the relationship between ungual bone geometry and certain lifestyles for a phylogenetically diverse sample of extant avians, this study overcomes limitations of past studies and enables direct comparison of fossil and extant material.

The claws of arboreal squamates had higher outer curvatures of digit III ungual bones compared to those of terrestrial squamates in the sample (Figure 4 A, B), which would suggest the results of this study represent a universal trend amongst tetrapods to some extent. However, our results indicate ungual bones of squamates have significantly lower curvatures than avian ungual bones and may be incomparable using these methods (Figure 4). It is unclear whether the relatively lower claw angles in squamates result from genetic signal or from alternate functional demands of quadrupedalism versus bipedalism influencing the evolution of claw morphology. As our group of squamates was very small (n=5) and including a phylogenetic outgroup would increase confidence of assertions for fossil taxa, further work including more outgroup taxa may be useful.

No significant correlation was found between claw angle and body mass for ungual bones. Only claw sheaths of terrestrial taxa exhibited a statistically significant relationship with body mass (Table 1). The relationship found is poorer than reported for some previous studies^8,70^ and roughly parallels findings of a recent comprehensive study^5^. These results suggest the correlations found in this study are relatively unaffected by body mass, and so the limited weight range of the extant sample taxa is not expected to have a significant impact on findings.

The predictive model based on ungual bone curvatures outperformed that based on claw sheath curvatures (Table 3), which suggests ungual bones provide a more accurate metric. This indicates reconstructing external claw morphology is unnecessary for comparative analysis. However, results based only on ungual bone morphologies alone were unstable and subject to change with the inclusion of sheath measures (Tables S1, S2). Models based on multiple measures taken on the ungual bone and on soft tissues of the claw yielded the most accurate predictions for extant taxa, and so when soft tissues are well-preserved in fossil claws we recommend following this approach.

Though median values for most behavioural categories could be separated based on one or more measures of claw curvature, there is significant overlap between the ellipses drawn for each behavioural category, particularly for that of claws belonging to perching birds. This results in frequent misclassifications of perching birds compared to other categories (Table 2). Predictions of lifestyle, particularly perching lifestyles, for fossil taxa based on claw morphology alone should thus be considered alongside additional evidence to improve reliability of predictions^71^.

The results in tables 5 and 6 show that reconstructing behaviour from structure is not particularly reliable. There are a number of reasons why this may be the case. Behavioural complexity presents an issue for this and any study attempting to link morphology with mode of life^5,9^. Most animals utilise pedal claws for multiple functions to varying degrees, and so it is difficult to class any animal into a single behavioural category^4,5,8^. Many birds with perching or climbing adaptations also spend time foraging on the ground, for example, and all predatory birds measured for this study are also perching birds. For this reason, one could alternately interpret the ‘predatory’ morphospace as a ‘predatory-perching’ morphospace. Unfortunately, it was not feasible to include ground-dwelling taxa that utilise claws for prey capture/dispatching, and so we cannot test if there is a distinction between claw morphologies of terrestrial versus arboreal predators.

Previous studies have found conflicting results regarding which, if any, of the defined groups exhibits greater behavioural generalization with regard to claw shape^5,8^. The results of this study suggest claw sheaths may more greatly reflect behavioural generalization or specialization, while ungual bones appear to possess roughly equivalent spread by group (Figure 4). Ungual bone curvatures of the predatory sample taxa have the narrowest morphospace and seem to have similarly-shaped claws. This may relate to biomechanics of piercing in prey capture and dispatching, or perhaps the predatory taxa could be interpreted as hyper-specialised perching birds. Alternatively, the narrow ‘predatory’ morphospace could be an artefact of the relatively small sample size for predatory claws.

In addition to the extant taxa measured, this study also measured 34 pedal claws from 26 specimens representing 18 genera of fossil dromaeosaurs, avialans, and anchiornithids. Our results suggest arboreal habits for many of the measured fossil taxa with the majority grouping with perching birds and roughly two-thirds plotting outside the 95% confidence ellipse for terrestrial taxa (Figure 4). Many fossil taxa that grouped with perching birds lack an opposable hallux and would have been incapable of perching in the style of modern birds. However, these results could be interpreted as supporting some degree of arboreality in Mesozoic theropods such as *Microraptor, Changyuraptor*, and *Pedopenna* that possess adaptations consistent with an arboreal lifestyle such as extensive hindlimb feathering^31,72,73^. Though true perching may have been impossible, it is plausible that curved pedal claws evolved to grip branches and/or tree trunks in conjunction with manus claws to enable arboreal behaviours.

*Archaeopteryx* claws were classed as either perching or predatory, and most received low posterior probabilities for a terrestrial classification (Table S1). The results could be interpreted as suggestive of arboreality and, more specifically, of a raptorial, perching lifestyle. However, *Archaeopteryx* claws were well scattered and recovered within shared morphospaces for perching, predatory, terrestrial, and scansorial taxa (Figure 4 C, D), and so *Archaeopteryx* cannot be assigned to any behavioural or ecological category with confidence based on these results.

One *Microraptor* claw classed as perching whilst another from the same specimen classed as predatory. Though conflicting predictions are of concern when inferring a specific mode of life, it is not unreasonable to conclude that sharply curved claws in taxa such as *Microraptor* were adapted for an arboreal lifestyle. This is supported by a fossil preserving an enantiornithine bird in the gut, which suggests the animal hunted in trees^75^, biomechanical models indicating gliding ability^73,75^, and certain anatomical features including hindlimb feathers that would have hampered terrestrial locomotion^74^. Of other Mesozoic theropods studied here, *Changyurapto*r and *Pedopenna* also seem likely candidates for arboreality based on their recovery within the overlap between predatory and perching morphospaces and various anatomical features consistent with arboreality^31,72^.

Multiple small dromaeosaurs fall within (*Microraptor, Changyuraptor*) or close to (*Zhenyuanlong*) the overlap between ‘predatory’ and ‘perching’ morphospaces, which suggests the foot may have been utilised in both prey capture and grasping branches in a niche analogous to that of modern raptorial birds. Multiple dromaeosaurs (*Halszkaraptor, Dakotaraptor*) plot outside the ‘predatory’ morphospace despite having carnivorous lifestyles, which supports our interpretation of the measured ‘predatory’ morphospace as, more specifically, ‘predatory-perching’. These large dromaeosaurs and one anchiornithid *Eosinopteryx* received robust terrestrial classifications (Figure 4 C, D; Table S1) consistent with osteological features and findings of past studies^36,45^; these predictions indicate phylogenetically high curvatures of paravian claws are not influencing false confirmations of arboreality as has been previously suggested^45^.

Interestingly, all claws belonging to *Confuciusornis* were classed as predatory by all models (Figure 4 C, D; Tables S1, S2). Conclusive direct evidence of diet is unknown for *Confuciusornis* specimens^76^, but beak morphology is traditionally regarded as evidence of herbivory in *Confuciusornis*^77^. However, our results suggest a predatory-perching ecology for *Confuciusornis* in conjunction with cranial morphology^78^ and the presence of an opposable hallux that would have enabled grasping of branches. Unfortunately, as some *Confuciusornis* claws fall outside of extant morphospaces (Figure 4 C, D), it is difficult to determine if the model’s predatory prediction is a robust inference of ecology. The predatory classification of *Anchiornis* (Table S1) is also suspect because all measured *Anchiornis* claws plot well outside extant morphospaces. These claws possess very high values for LD2 because the ventral arcs of the claws have significantly greater angles of curvature than the dorsal arcs of the claws. High LD2 values are used to distinguish the predatory morphospace, and so the model classes these fossil claws as predatory despite data points falling outside extant morphospaces.

Predictions of predatory ecology for two *Jeholorni*s claws contradict direct evidence of granivory for at least five specimens preserving seeds in the gut^76^. One claw received a perching prediction and so it seems probable based on this, direct dietary evidence, and skeletal characters that Jeholornis was a non-raptorial perching bird that the model struggled to classify. Claws from *Sapeornis* classed as either perching or scansorial, either of which could be argued for the taxa based on skeletal characters^79^. Other early avians *Zhouornis, Eopengornis, Parapengornis*, and *Fortunguavis* received predictions of terrestrial and perching but generally could not be well-resolved. Early birds are believed to have been well-diversified by the late Mesozoic^80^, and so it is possible that terrestrial and perching lifestyles are represented in the dataset. However, *Zhouornis* received conflicting predictions for claws of the same specimen, and the perching prediction for *Fortunguavis* contradicts interpretations of a scansorial lifestyle based on skeletal characters^34^.

If we are to take these results at face value then it paints an interesting picture of the evolution of arboreality in the avian lineage. Though classifications varied widely and often conflicted for claws of the same genus or even from the same specimen, the results suggest at least some of fossil paravians engaged in an arboreal lifestyle. The prevalence of arboreal predictions for fossil paravians and early avialans suggests arboreal behaviours evolved in some ancestor of birds prior to the origination of Aves. This in turns supports the “trees-down” hypothesis of powered flight evolution in modern birds.

The models occasionally yielded conflicting predictions for left and right digit III claws belonging to a single fossil specimen. This could be caused by natural variation within the population, taphonomic distortion, or some unknown factor. The measured fossil claws did not exhibit any obvious distortion and often plotted within the overlap between morphospaces, and so it is plausible slight, naturally-occurring differences between claw angles could result in conflicting predictions. Conflicting predictions also occurred between specimens for *Jeholornis, Sapeornis*, and *Archaeopteryx*. In addition to previously suggested causes, it is possible these conflicting predictions occurred because the specimens represent different species with different modes of life. Regardless of causative factors, conflicting predictions hinder classification of fossil taxa and should be considered in any future work seeking to class fossil taxa based on claws.

## CONCLUSIONS

Analysing ungual bone morphology has clear benefits in palaeontological study over the analysis of external claw morphology. The study found that curvatures of the ungual bone not only provide a useful proxy for certain modes of life but in fact exhibit a stronger correlation with lifestyle (kappa=0.7141) than do similar measures taken on claw sheaths (kappa=0.6324). However, utilising solely curvatures of the pedal digit III ungual bone to predict lifestyle is ill-advised because morphospaces overlap, and it is difficult to determine if fossil and extant claws are truly comparable based on these results. We suggest curvatures of fossil ungual bones may be useful when studying fossil taxa as they exhibit a strong correlation with behaviour and ecology for modern birds and squamates, but other osteological features and phylogeny should be considered.

## Acknowledgements

This work was supported by funding from NERC (grant number NE/R011168/1). The authors thank The World Museum of Liverpool, The Manchester Museum, and The National Museum of Scotland at Edinburgh for their contribution of specimens for this study. The authors also thank Dr. Andrew Chamberlain for his support with X-ray equipment and Callum McLean for manuscript revisions.

